# Effect of Hydromethanolic Seed Extract of Garcinia Kola on Male Reproductive Hormones

**DOI:** 10.1101/2022.12.14.520482

**Authors:** Tamunotonye Harry

## Abstract

Reproduction is central to the continued existence of mankind on earth, while infertility presents both a social and a public health concern, with male factor infertility present in 20% – 50% of all cases of infertility among couples. As part of the solution for male fertility disorders, drugs including those that are plant-based are being employed to address such issues. Hence, this study was focused on the effect of the administration of hydro-methanol seed extract of *G. kola* on the reproductive hormone of adult male Wistar rats. The plant material was procured, identified, and extracted while a total of 36 adult male rats were procured and distributed equally into three groups which were subdivided into two. Animals in group 1 served as the control while groups 2 and 3 served as the extract-treated groups which received 100 and 200 mg/kg BW of the extract respectively. Those in subgroups A and B were treated for 14 and 30 days respectively after which the animals were sacrificed and the blood collected via cardiac puncture into the appropriate bottles for hormonal profile assay. The result showed that the administration of the extract resulted in a reduction in the level of FSH and LH but increased the level of Testosterone, with a significant increase recorded in the 200 mg/kg group, while the level of prolactin was increased in the 100 mg/kg group but decreased in the 200mg/kg group. Hence, Garcinia kola might not be the best solution to aid fertility in males

## INTRODUCTION

Reproduction is central to the continued existence of mankind on earth, while infertility which is a major problem associated with reproductive health in the world has grown to the extent that it is now a social concern and a public health concern (Anate et al., 2002). Infertility may be defined as a biological inability to achieve conception after one year of unprotected coital exposure. Globally, about 50 - 80 million people suffer from infertility, with an estimate from the World Health Organization (WHO) showing that approximately 8 - 10% of couples suffer from this problem, while the male factor infertility is present in 20% – 50% of these couples, either independently or in conjunction with female factor infertility issues (Jarow et al., 2002, Singh et al., 2014). In the context of males, a man is said to be infertile if he is unable to impregnate his partner after one year of unprotected intercourse (Singh et al., 2014). This condition may result from a number of factors including hormonal imbalances, genetic defects, chemicals, and toxins exposure reproductive anatomical and morphological abnormalities, reactive oxygen species, and smoking (Aboua et al., 2013; Singh et al., 2014).

As part of the solution for male fertility disorders, drugs including those of plant-based are being employed to address such issues. Historically, plants (including their components) have provided a source of inspiration for novel drug compounds, as plant-derived medicines have made large contributions to human health and well-being. One such plant is *Garcinia kola* which has a bitter astringent and resinous taste, somewhat resembling that of raw coffee, followed by a slight sweetness, hence it is commonly known as bitter kola. Bitter cola is a highly valued ingredient in African ethnomedicine because of its varied and numerous uses which are social and medicinal. It is also chewed extensively in Southern Nigeria as a masticatory to cause nervous alertness and has been proven to exhibit pharmacological uses in treating coughs, throat infections and liver disorders due to its phytochemical content (Farombi et al., 2005; Ogunmoyole et al., 2012). According to Akpantah (2003), *Garcinia kola* has been reported to increase spermatogenic activity through its tissue enhancement and ability to increase peripheral testosterone, while Ralebona et al., (2012) claimed that the plant possesses aphrodisiac effects and as such is used traditionally in the treatment of erectile dysfunction. These reproductive properties can be linked to the phytochemical contents of the seed which as discovered by Essien and Effiong (2014) include tannins, saponins, flavonoids, alkaloids, and cardiac glycosides, while Ogunmoyole et al (2012) reported the presence of antioxidants such as total phenolics, flavonoids, and vitamin C content. Hence, the focus of this study was to determine the effect of the administration of hydro-methanol seed extract of *G. kola* on the reproductive hormone of adult male Wistar rats.

## MATERIALS AND METHODS

### Plant Material and Extraction

The *Garcinia kola* seed used for this study were bought from Choba market, Rivers state and subsequently identified with Herbarium number UPH/V/1220, by the taxonomist in the Department of Plant Science and Biotechnology, University of Port Harcourt, Nigeria. Voucher specimen of the plant was deposited in the herbarium. The seeds were dried and the covers removed before they were ground into powder prior to its extraction at the Pharmacognosy department of the University of Port Harcourt, using Soxhlet extractor, with hydromethanol (80% methanol) as solvent. The solution was filtered using Whatman filter paper and the filtrate concentrated under reduced pressure in vacuum at 45°C. The extract yield was transferred to a hot oven where they were dried to a constant weight at 45°C and stored at 4°C. The extract was resuspended in distilled water before administration.

### Experimental Animal

A total of 36 adult wistar rats were procured from the animal house of Department of Human Physiology, University of Port Harcourt and used as the experimental model. The animals which had an average weight of 100g, were handled under the laboratory conditions, in accordance to National and Institutional guidelines for animal usage in experimental purposes and fed with standard feeds and water. The animals were also allowed to acclimatize for 14 days before the experimental processes began.

### Experimental Design

The 36 male wistar rats recruited for this study was divided into 3 groups of 18 animals each. The animals in Group 1 served as the Control group and receive 1ml distilled water while the animals in Groups 2 and 3 served as the treatment group and received 100 and 200 mg/kg BW of hydromethanol seed extract of *G. kola* respectively. The animals in each of the groups were further divided into two whereby 6 rats in each of the groups were treated for 14 days before sacrifice while the second 6 rats were treated for complete 30 days before sacrifice. The entire administration was done orally once daily throughout the duration of the administration, while the dosage of the extract administered to the animals was below the mean lethal dose (1000 mg/kg) as reported by Essien and Effiong (2014).

At the end of the administration periods for each of the sub-groups, the animals were sacrificed under light chloroform anesthesia. The blood was also collected via cardiac puncture into the appropriate bottles for hormonal profile assay. The sera samples were separated and then assayed for Testosterone, FSH and LH using enzyme-linked immune-absorbent assay (ELISA) as described in the study of Akighir, Inalegwu and Onyezili (2018).

### Statistical Analysis

Data were statistically analyzed using SPSS version 21 and the results expressed as mean ± standard error of mean (SEM). Significant differences were determined using one-way analysis of variance (ANOVA), while a p-value of less than 0.05 (*p<0*.*05*) was considered statistically significant.

## RESULTS

### Male Reproductive Hormone Profile

The result of the effect of the administration of the hydromethanol seed extract of *G. kola* on the male reproductive hormones is presented in Table 1 above. According to the finding, 14 days administration of the extract at both doses did not produce any significant change in the FSH levels (3.44±0.14 and 3.43±0.6) in comparison with the control group (3.42±0.12), while the LH level in both treatment groups (4.84±0.07 and 4.89±0.28) were non-significantly (p>0.05) lower in comparison with the control (5.01±0.32). On the other hand, the Testosterone levels in both treatment groups (2.36±0.27 and 1.01±0.43) were significantly (p<0.05) lower in comparison with the control group (3.82±1.01), while the extract did not produce any statistically significant difference with the level of the prolactin (4.20±0.17 and 4.30±0.37) in comparison with the control group (4.19±0.09).

**Table 1:** Effect of hydromethanol seed extract of G. kola on hormone profile of wistar rats.

| Groups | 14 days treatment |  |  |  | 30 days treatment |  |  |  |
| --- | --- | --- | --- | --- | --- | --- | --- | --- |
| | FSH<br>(m/$\mu$/L) | LH<br>(m/$\mu$/L) | T<br>(ng/μL) | P<br>(ng/μL) | FSH<br>(m/$\mu$/L) | LH<br>(m/$\mu$/L) | T<br>(ng/μL) | P<br>(ng/μL) |
| **Control** | 3.42±0.12 | 5.01±0.32 | 3.82±1.01 | 4.19±0.09 | 0.38±0.14 | 14.28±9.84 | 2.80±1.48 | 0.79±0.47 |
| **100**<br>**mg/kg**<br>**Extract** | 3.44±0.14 | 4.84±0.07 | 2.36±0.27* | 4.20±0.17 | 1.20±0.64 | 4.38±3.63 | 4.40±2.88 | 6.50±7.59 |
| **200**<br>**mg/kg**<br>**Extract** | 3.43±0.6 | 4.89±0.28 | 1.01±0.43* | 4.30±0.37 | 2.58±1.03* | 2.00±2.19* | 3.00±0.81 | 0.50±0.18 |

Data are expressed as mean ± SEM, n=6
\* = significantly different from control (P<0.05)
FSH = Follicle Stimulating Hormone; LH = Luteinizing hormone; T = Testosterone; P = prolactin

After prolonged administration of the extract for 30 days, result showed that the level of FHS in the 200 mg/kg extract treated group (2.58±1.03) was significantly (p<0.05) higher in comparison with the control group (0.38±0.14) while that of the 100 mg/kg extract treated group (1.20±0.64) produced non-significant (p>0.05) increase in comparison with the control group. The level of the LH followed an opposite trend which showed that the 200 mg/kg extract treated group (2.00±2.19) was significantly (p<0.05) lower in comparison with the control group (14.28±9.84) while that of the 100 mg/kg extract treated group (4.38±3.63) produced non-significant (p>0.05) lower in comparison with the control group. On the other hand, the prolonged administration of the extract did not produce any significant change in the level of the prolactin and Testosterone when compared with that of the control. However, the level of testosterone in both treatment groups (4.40±2.88 and 3.00±0.81) were higher in comparison with the control (2.80±1.48), while the level of the prolactin in the 100 mg/kg extract treated group (6.50±7.59) was higher in comparison with the 200 mg/kg extract treated group (0.50±0.18) and the control group (0.79±0.47).

The study also analysed the relative percentage change of the male reproductive hormones after the 14 days and 30 days administration period with the result presented as figure 1 below. According to the finding, the FSH level in both the 100 and 200 mg/kg groups decreased by 65.1% and 24.8% respectively, while that of the LH decreased by 9.5% and 59.1% respectively. On the other hand the level of testosterone was seen to increase by 85.6% and 200% in the 100 and 200 mg/kg extract treated groups while the level of prolactin in the 100 mg/kg group increased by 54.8% and that of the 200 mg/kg group decreased by 88.4%

**Figure 1:**
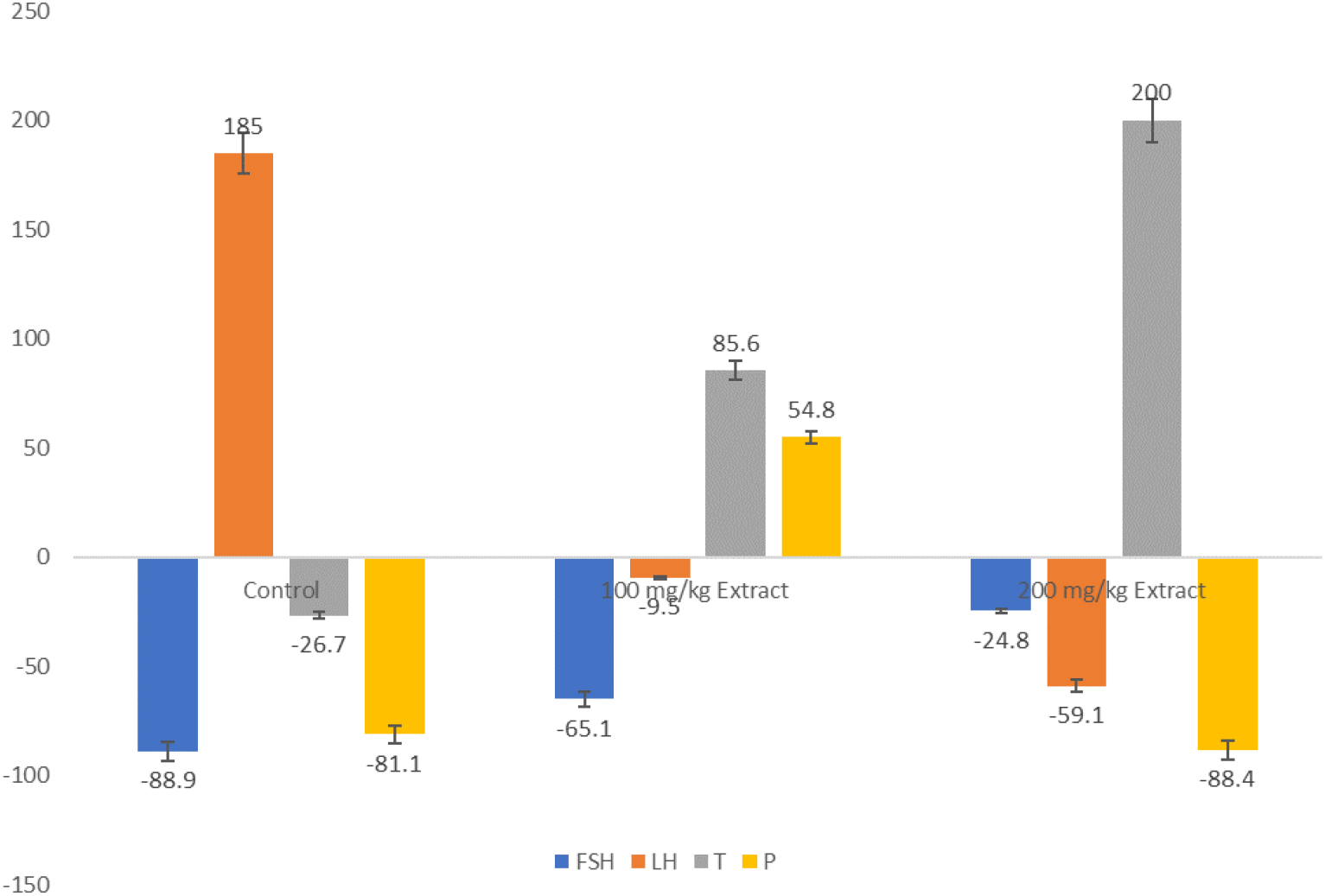
Relative change (%) of the effect of hydromethanol seed extract of G. kola on hormone profile of wistar rats after 14 days and 30 days of administration.

## DISCUSSION

According to Iyke et al. (2018), FSH play vital roles in gonadal development, steroidogenesis and maturation at puberty as well as gamete production (spermatogenesis) during fertile life, while the LH is known to stimulate the secretion of sex steroids (Testosterone) from the gonads. As seen in this present study, the administration of the hydromethanol seed extract of *G. kola* caused a reduction in the level of the FSH and LH but increased the level of Testosterone, with a significant increase recorded in the 200 mg/kg group. The 100 mg/kg administration of the extract caused an increase in the level of Prolactin while the 200 mg/kg administration caused a reduction. This is similar to the findings of Ralebona et al., (2012) where different doses of ethanolic extract of G. kola seeds increased the Testosterone levels, while Akpantah (2003) reported that the increased spermatogenic activity as seen in their study could be linked to increase peripheral testosterone. An earlier study by Braid et al., (2003) presented a contrary report which showed that the administration of methanolic alkaloid extract of *Garcinia kola* seed caused marked reduction in serum testosterone and a concomitant elevation of serum FSH and LH in males, while Yakubu and Quadri (2012) reported that the serum testosterone, FSH and LH were not significantly altered by the administration of the aqueous seed extract of *G. kola*. Eyong and Braide (2009) also reported that prolonged (7 weeks) oral administration of the alkaloid extract of *Garcinia kola* seed resulted in a significant elevation of serum concentrations of LH and FSH, but a reduction in serum levels of testosterone, while Agube (2001) also reported that the activity of *G. kola* may influence reproductive activities through its action on gonadotrophins and testosterone by reducing the serum level of testosterone and markedly elevating the levels of FSH and LH in male rats. Also, Obiandu and Adienbo (2018) discovered that the administration of the pulp extract of Garcinia kola elevated the level of the LH and FSH, while in line with this study, the Testosterone level was found to be significantly increased with the higher dose of the extract. These differences could be due to differences in the study designs, the dose of administration as well as the duration of the administration in the different studies. However, failure of the pituitary gland to secrete the gonadotrophin hormones (FSH and LH) as seen in this study, could lead to a disruption of testicular function and thereby result to infertility in males (Obiandu &Adienbo, 2018). additionally, Gelain et al. (2005) reported that decreased levels of FSH is implicated in reduced level of spermatogenic activities, occurrence of infertility and reproductive toxicity. Hence, in line with the suggestion of Braid et al., (2003), the extract possesses possible antifertility consequences.

## CONCLUSION

This study which was designed to examine the administration of hydromethanol seed extract of *G. kola* on the male reproductive hormone was able to show that the extract reduced the level of gonadotropins (LH and FSH) but increased that of the Testosterone in both doses, while prolactin was increased only in the low dose. In the light of this study and its findings, Garcinia kola might not be the best solution to aid fertility in males and so we recommend that people in sub-saharan Africa (SSA) and especially in Nigeria reduce their intake of bitter kola seed.

